# NAMERS: a purpose-built reference DNA sequence database to support applied eDNA metabarcoding

**DOI:** 10.1101/2023.10.06.561210

**Authors:** Kristen M. Westfall, Gregory A. C. Singer, Muneesh Kaushal, Scott R. Gilmore, Nicole Fahner, Mehrdad Hajibabaei, Cathryn L. Abbott

## Abstract

Applied eDNA metabarcoding is increasingly being used to generate actionable results to inform management decisions, regulations, or policy development. Because of these important downstream considerations, optimizing workflow elements is now essential to increasing standardization, efficiency, and confidence of metabarcoding results. Reference DNA sequences are critical workflow elements that currently lack consistent approaches to generating, curating, or publishing. Here we present a complete (mitochondrial genome and nuclear ribosomal DNA cistron) and high quality reference DNA sequence library for the freshwater fishes of British Columbia, Canada. This resource is published as the Novel Applied eDNA Metabarcoding Reference Sequences (NAMERS) repository (https://namers.ca), a user-friendly and interactive website for specialists and non-specialists alike to explore and generate custom reference libraries for taxa and genes of interest. We demonstrate the power of NAMERS to optimize applied eDNA metabarcoding workflows at the study design stage by analyzing the number of primer mismatches and resolution power of existing metabarcoding markers. To meet the increasing demand for actionable eDNA metabarcoding applications, NAMERS demonstrates that high quality curated genomic information is within a reasonable reach. It is timely to establish this framework as the new gold standard and coordinate our efforts to generate this type of reference data at scale.

## INTRODUCTION

The need has never been greater for information rich biomonitoring data to assess environmental impacts, monitor rare species, chart ecosystem trajectories, and evaluate remediation and conservation efforts - all to ultimately maximize positive desired outcomes. There is high current attention on translating the science of eDNA metabarcoding into practices that benefit humankind ^1–7^. Despite the proven usefulness of eDNA metabarcoding for biomonitoring, widespread uptake for decision-making requires additional attention to workflow elements that are essential for translating the science to a broad end-user community. Specifically, optimizing factors related to operational feasibility, cost, taxonomic breadth, throughput, and scalability, as overviewed in other works^1,3,8^. Many sources can introduce potential biases in the metabarcoding workflow^9^. Indeed, the lack of quality criteria or standards and unknown confidence in results is a serious impediment to the uptake of eDNA methods for applications^10^.

Here we adopt the term *applied eDNA metabarcoding* to distinguish between scenarios where eDNA results are generated for practical goals (i.e. inform a management decision) versus foundational research (i.e. to advance basic knowledge). This distinction is important for recognizing the quality and confidence requirements needed for eDNA results to become accessible to end-users to inform decision-making, including for building end-user trust^11^. Darling et al.^12^ argued for the need to distinguish between high throughput sequencing studies designed for ecological versus biosecurity surveillance purposes for the same reason and the term applied eDNA metabarcoding brings this idea a step further to encompass all practical uses.

DNA barcoding emerged as the gold standard for genetic species identification before the naissance of eDNA metabarcoding almost a decade later^13,14^. DNA barcoding uses a standardized DNA sequence proven to provide species-level resolution and a purpose-built, curated database (the Barcode of Life Data System^15^) to determine the taxonomy of unknowns. In contrast, eDNA metabarcoding often uses multiple markers not proven to amplify all target taxa or to provide species level resolution therein, and assigns taxonomy to unknowns using data from massive online global DNA sequence repositories^1,6^ (e.g GenBank^16^). Not only do these repositories generally lack curation and rigorous quality control, but they are often amplicon based, meaning primer region sequences cannot be assessed for complementarity to target taxa and novel primer design to meet specific needs is precluded.

It has been known since the inception of eDNA metabarcoding that standardized DNA barcoding markers will not suffice given the need for marker choice flexibility and that comprehensive databases of whole organellular genomes are required^14^. Regardless, most references to DNA sequence databases for eDNA metabarcoding focus on filling gaps in existing single gene repositories^17,18^. There has been a lack of recognition of the *need for innovation around reference DNA sequence databases* to meet the unique needs of applied eDNA metabarcoding. This is despite it being well proven that incomplete or low quality reference data can lead to biased or erroneous results, including false positives and false negatives^9,19,20^ that quickly erode the confidence of potential end-users. Sound reference data are essential for both the front and back ends of the eDNA metabarcoding workflow: they are vital for robust survey design, as related to primer design and/or marker selection, and for interpretation of metabarcode sequences, as related to taxonomy assignment.

Here we present the development of a mitogenome (and ribosomal DNA cistron) web portal as a new approach, with these as basic premises of applied eDNA metabarcoding: (i) primer design is a key factor determining success^21,22^, and primers must be chosen carefully to meet application-specific needs^14,23^; (ii) achieving species resolution is important, as higher taxonomic levels lack sufficient information for most biomonitoring needs^8^, (iii) different genes work best for different taxa^21,24^, and multi-marker methods out-perform single marker ones if reference databases are sufficiently populated^25^; and (iv) mitochondrial markers are the workhorses of vertebrate eDNA metabarcoding with gene regions other than COI working best^23,26^.

On this basis we developed NAMERS: <u>N</u>ovel <u>A</u>pplied eDNA <u>Me</u>tabarcoding <u>R</u>eference <u>S</u>equence database, a web portal of whole mitogenomes and nuclear ribosomal (nr) DNA cistrons for freshwater fish in British Columbia, Canada, with specific functionalities designed explicitly for applied eDNA metabarcoding. This database improves DNA reference sequence data quality, completeness, and accessibility, as well as transparency to end-users, and can be geographically and taxonomically expanded over time to include all Canadian freshwater species. We demonstrate several important benefits of NAMERS for applied eDNA metabarcoding compared to existing massive global DNA sequence repositories, including: increased end-user confidence given quality control measures; even coverage of all mitochondrial genes; easy construction of complete custom libraries for any gene; and greatly improved time-efficiency and user-friendliness for specialists and end-users alike. The portal offers ‘one-stop shop’ access to instant downloads of sequence alignments and complete reference libraries from high quality data, requiring no bioinformatics experience. While mitogenome sequencing may seem an intimidating task, if one accepts the premises of applied eDNA metabarcoding presented above, it is starkly clear that gap-filling single gene repositories does not meet the mission-critical needs of this field. Thus the added investment in mitogenomes is not only well justified but essential.

## RESULTS

### Shotgun DNA sequencing, assemblies, and annotation

Mitogenomic data were generated here for an estimated 92.3% (85/92) of all freshwater fish taxa present in our target geographic area of BC, Canada, representing 49 genera and 19 families. Sequencing success rates were high; 82 of 85 taxa sequenced returned complete or near complete (missing one gene or few partial genes) mitogenomes and a further three returned partial mitogenomes. Thus the final data set is comprised of complete or near complete mitogenome sequences for ~89% of all freshwater fish taxa in BC (82/92) and partial mitogenomes for an additional two species and one lineage. Mitogenome sequencing depth ranged from 1.4 to 2249.9 with a median of 101.2, and mitogenome length ranged from 14.198 – 18.142 kbp with a median of 16.625 kbp. All species for which the full mitogenome was constructed contained 13 protein-coding genes (COX1 – COX3, CYTB, ND1 – ND6, ND4L, ATP6, and ATP8), 22 tRNA genes, and two rRNA genes (small and large rRNA subunits). Full nrDNA cistrons containing 5.8S, 18S, and 28S regions were sequenced for 70 species, with sequencing depth ranging from 16.5 to 1340.8 with a median of 413.4. Full details on mitogenome and nrDNA data are in Supplementary Tables 1 to 3.

### Quality assurance

The morphological taxonomy of each specimen in NAMERS with genetic identification by COI barcoding in almost all instances, with a few exceptions detailed here. The NAMERS Umatilla dace (Cyprinidae; *Rhinichthys umatilla*) specimen was a misidentified sucker (Family Catostomidae), which is highly plausible given the difficulty identifying juvenile fish, and hence excluded. For lampreys, both the COI and 12S genes were not variable within genera but the 16S gene differentiated the *Entosphenus* species and the cytochrome b (CYTB) gene differentiated the *Lampetra* species (excluding the Morrison creek variant), by a single base in all cases. The ND4 gene had greatest genetic variation with three base pair differences between the two *Entosphenus* species and two differences between *Lampetra* species (again excluding the Morrison Creek variant); thus ND4 would be the best candidate gene for species specific markers in this family.

### NAMERS portal specifications

Whole mitogenomes, annotated mitochondrial genes, and annotated nuclear ribosomal genes are available to view and download in FASTA format on the new NAMERS portal. The main database page offers a table of 86 species grouped by increasing taxonomic levels. Users can highlight any taxonomic level to view available sequence data for all included taxa, from individual species to family, and can easily customize download batches of particular genes or taxa. They can also highlight particular genes of interest or the complete mitogenome for automatic alignments (using MUSCLE^27^) of selected genes and taxa of interest; the alignment will automatically update when the user engages or disengages genes or taxa. Clicking on an individual species will take users to a species page containing specimen metadata, mtDNA and nrDNA *de novo* assembly statistics, mitogenome circular sketch map, and links to download the full mitogenome or annotated genes in FASTA format.

The number of primer mismatches and the proportion of species resolved for 19 published fish and vertebrate metabarcoding markers was assessed using the NAMERS database (Table 1 and Figure 1). Due to low genetic variation already reported within Petromyzontidae, no metabarcoding marker could differentiate species (Figure 2), therefore we report species resolution rates with and without lamprey included (Figure 1). Across all 86 taxa (representing 19 families), the number of mismatches in forward and reverse primers was highly variable, ranging from 0 to 8, with 12S and 16S primers generally having fewer than COI and CYTB (Figure 1). The single marker with the fewest overall mismatches was Vert16S^28^. Excluding lampreys, species resolution rates varied from 75% to 100% (Figure 1).

**Table 1:**
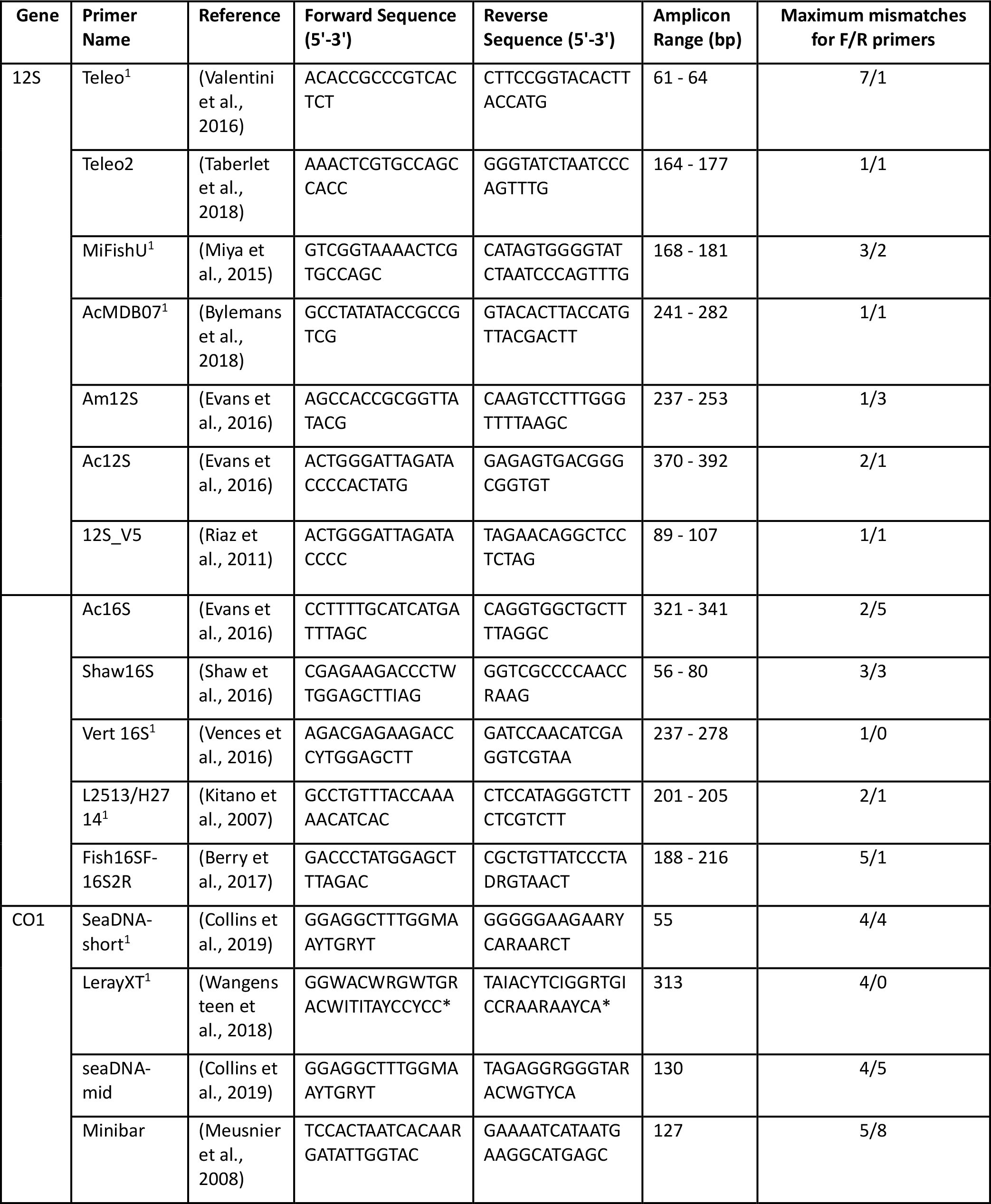
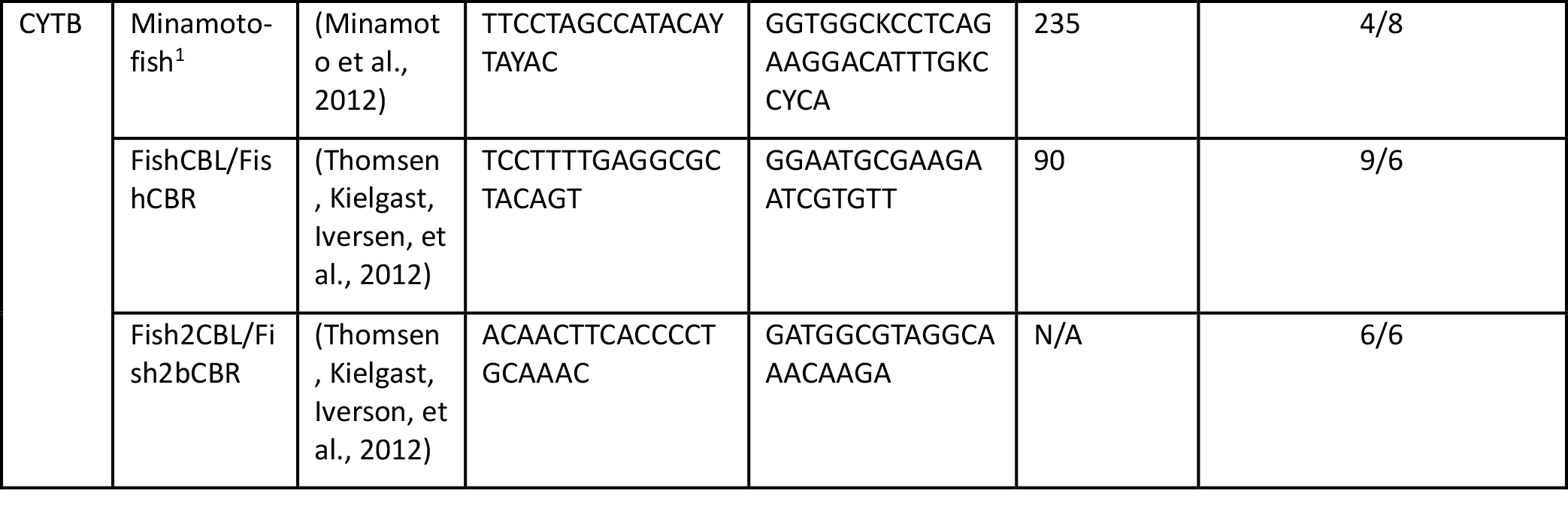
Information on markers assessed for primer mismatches and species level resolution in 82 freshwater fish species. ^1^Markers presented at the Family level in Figure 2.

**Figure 1:**
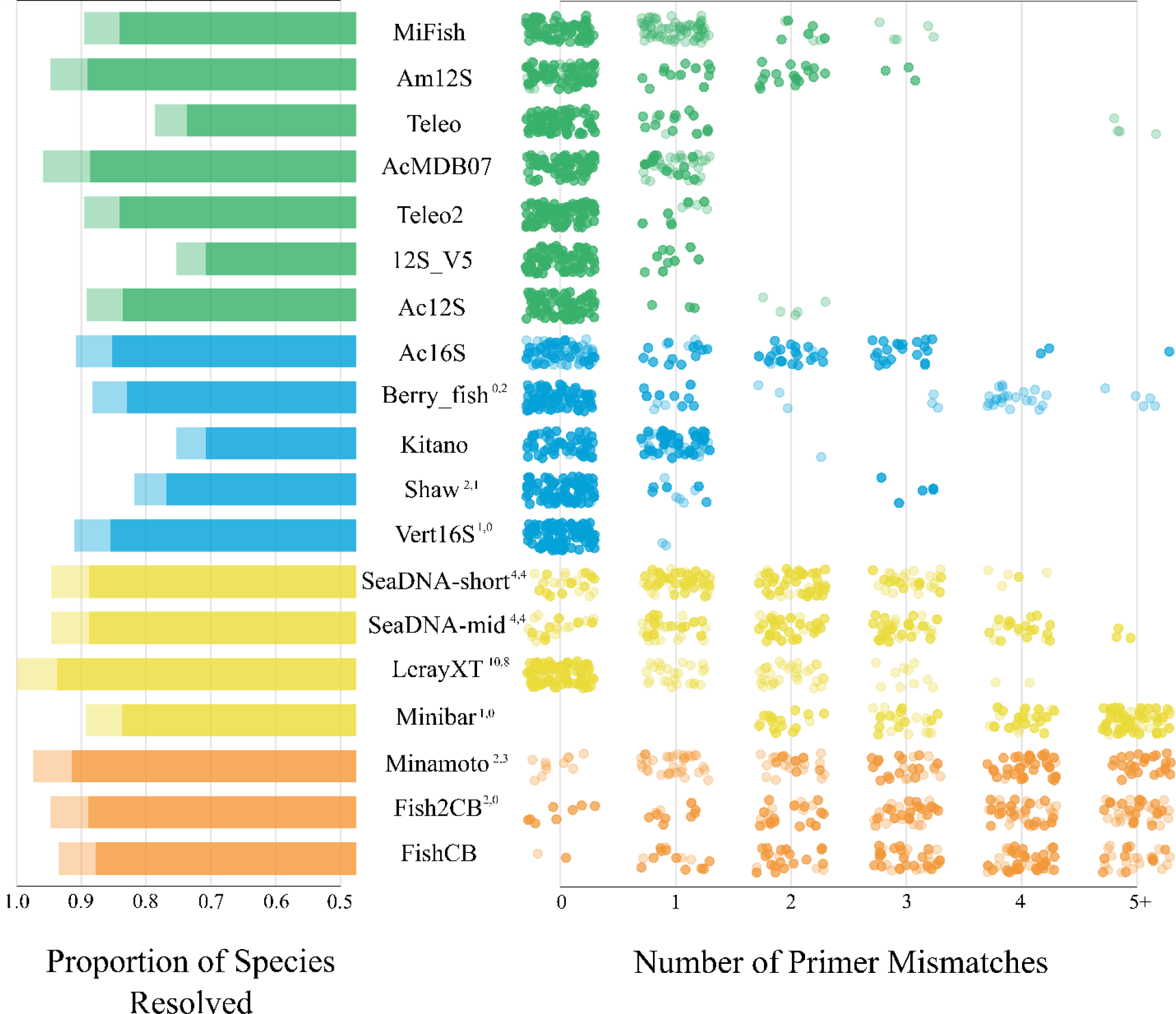
Primer mismatch and species resolution data for 19 markers from four gene regions, generated using all species in the NAMERS database (n=86). On the right hand panel is the number of mismatches between primer and species from zero to four and then 5 and over (5+, the maximum was 9). Forward primer mismatches are depicted by dark circles and reverse primer mismatches by light circles. On the left hand panel is the proportion of species resolved by each marker (i.e. distinguished by a minimum of one base pair including indels); the dark coloured bar area depicts resolution when lamprey are included and the entire bar depicts resolution when lamprey are excluded. Superscripts indicate the number of ambiguous bases in the forward and reverse primers, respectively. Markers with no superscripts have no ambiguous bases in either primer.

**Figure 2:**
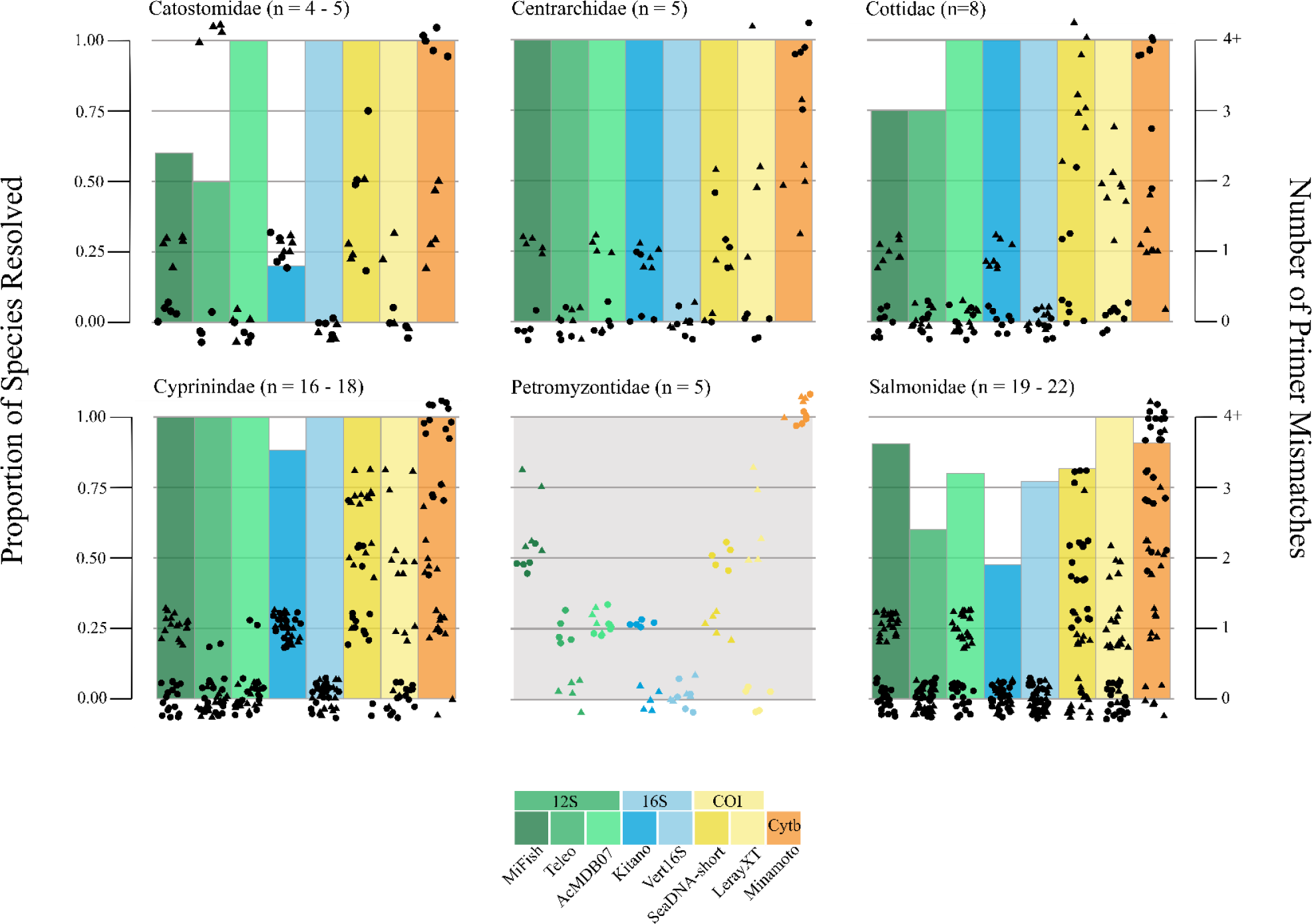
Family level plots of the number of primer mismatches, depicted by solid black symbols and the right-hand y-axis (forward primer = 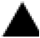, reverse primer = 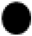), and the proportion of species resolved by unique amplicons defined as a minimum of 1 bp difference including indels, depicted by coloured bars and the left hand y-axis. Plots are for all families in the NAMERS database with a minimum of five species and a subset of eight of the primers tested in Figure 1. Note that no species in Petromyzontidae were resolved by any marker. The range in number of species per family used in the analysis reflects missing data; some species had some missing or partial genes available for analysis (See Supplementary Tables 2 and 3).

Achieving species level resolution within families will at times be more important than doing so across all families, as some applied eDNA metabarcoding surveys will target lower taxonomic groups only to ensure robust performance for the specific survey aim. The number of primer mismatches and proportion of species resolved at the family level are shown in Figure 2 for families (n=6) with more than five species (up to 63 species). Species resolution rates within families ranged markedly from none in lampreys (Petromyzontidae, n=5) to 100% in sunfishes (Centrarchidae, n=5) across all markers tested (Figure 2). Noteworthy is that in four of the six families in NAMERS with at least five species, Vert16S resolved 100% of species with zero primer mismatches (Figure 2).

## DISCUSSION

We are facing unprecedented rates of global change and biodiversity loss, and applied eDNA metabarcoding has the potential to augment the information content of current biomonitoring programs and elevate our capacity to successfully manage a changing biosphere. It is now accepted that minimum quality criteria and communicating sources of uncertainty are vital for translating eDNA based monitoring results into sound decision-making^10,20,29,30^. Here we introduce the term applied eDNA metabarcoding to facilitate the recognition of the unique quality- and confidence-related needs that come with translating this eDNA method into practical application. We suggest this translation is dependent on two key areas within the domain of science itself (i.e. there are other relevant factors that are cross-sectorial, e.g. communication between scientists and end-users): (i) standardization and quality control, to increase confidence in results^14,18,31^; and (ii) accessibility and transparency of methods and data^7^.

Reference DNA sequences are a challenging element of the eDNA metabarcoding workflow for which to satisfy these two key areas. Global genetic repositories do not have the level of curation necessary for defensible species assignments from environmental sequences^31^, and are problematic with respect to low quality control, reproducibility, transferability, and ease of regular updating^32^. Whilst some level of error correction is achievable^32^, several elements that lead to errors cannot be corrected, including uneven taxonomic coverage, undocumented specimen metadata, and lack of voucher availability or quality control processes. Further, since eDNA metabarcoding tools will often be multi-marker and implemented at local and regional scales, rarely global ones, the use of both single gene repositories and massive sequence databases is impractical as generating custom libraries from these is extremely laborious and requires specialized expertise. Without a dedicated quality-controlled repository for applied eDNA metabarcoding reference sequences, there is no means of preventing even high quality DNA sequence data from ending up in the abyss of large repositories where it is intractable to filter from the rest.

The need for improved reference data is repeatedly highlighted in studies^33^ yet there is no consensus on what the goal should be (i.e., sets of genes vs mitogenomes). Here, NAMERS offers multiple functionalities and access to mitogenome and rDNA sequences from vouchered specimens with the same level of associated metadata. It is NAMERS user friendly and publicly accessible, whereby users can instantly view multiple species alignments for taxa and genes of interest, download custom reference sequence libraries, and find links to each species’ record in a leading global biodiversity information system for finfishes (FishBase : A Global Information System on Fishes^34^). The ability to easily view and download a custom reference library of complete mitochondrial and nrDNA genes in NAMERS brings this type of repository into a new era with functionalities that set it apart from existing mitogenome databases. Furthermore, the NAMERS database is versioned to control for changes in taxonomic classification and additional taxa or geographic representatives. This increases transparency in data provenance for end users with respect to any changes to the database and their impact on the interpretation of metabarcoding results.

Species richness can be underestimated by indiscriminate application of metabarcoding markers without a full understanding of their specificity for target groups or level of species coverage within the reference library used for taxonomy assignments^35^. It is vital to make informed decisions on marker choice at the outset of each application. NAMERS enables this given it’s high taxonomic completeness of 94% partial and 89% full mitogenomes of all described freshwater fish taxa in BC (these numbers will be improved to 100% over time). We also demonstrate here the integral role of complete mitogenomes for defensible marker selection based on species resolution levels and primer complementarity. This is vital for ensuring accuracy in general or taxon-specific applied eDNA metabarcoding studies and yet is largely ignored in the literature and not possible using amplicon libraries for reference sequence data. Furthermore, amplicon libraries perpetuate the use of a narrow range of markers based on their use in previous studies, despite that marker choice needs to be carefully made to ensure fitness-for-purpose.

A large range in primer mismatches was identified across all species and markers tested here, but general patterns emerged. Although average species resolution was greater for the protein coding COI and CYTB genes as expected, the small and large ribosomal subunits had lower average rates of primer mismatch and would therefore likely recover more freshwater fish species when conducting taxonomically broad surveys. In some instances, family- or genus-level species resolution may be the priority, such as for conservation or identification of invasive species^36^. These specific applied eDNA metabarcoding end-user needs are common at multiple levels of government yet are perhaps somewhat less represented in the primary literature. Taxonomically complete reference libraries such as NAMERS enable in-depth analyses of family level resolution; we showed that variation in species resolution rates among families informs marker choice differently than overall species resolution rates across families. For example, no marker could identify any of the five lamprey species (Petromyzontidae, Figure 2) present in BC, but we identified variation in the ND4 gene that could potentially be used to design new markers for this family. Salmonids are a culturally and economically important species in BC, and only one marker could differentiate all species. Although the LerayXT marker had up to four mismatches across all families (Figure 1 and Table 1), we find only a maximum of two mismatches with the Salmonids (Figure 2), making it a viable marker for studies focusing on this family.

It is no longer far-fetched to make whole mitogenomes the new standard for reference DNA sequences given genome skimming capabilities^37^ and that the cost-per-base for ultra high throughput sequencing (i.e., NovaSeq) steadily decreases. The NAMERS repository is presented here as a framework or foundation to inspire this investment and motivate the considerable coordination that will be necessary to extend it to include larger geographic and taxonomic scales. The NAMERS database echoes Dziedzic et al.^17^ in demonstrating that a small research group can generate a comprehensive set of high quality mitogenome reference sequences for a geographically relevant area, which can be impactful on it’s own or within a larger framework. We acknowledge that long term support and continued augmentation of any database like NAMERS requires dedicated resources, and yet argue the need for this and the plausibility of this given the increased need for metabarcoding for real-world applications. While there are inherent limitations to any diagnostic approach especially for large-scale and taxonomically broad biota, eDNA metabarcoding has proven “good enough” for a large number of use cases^38^. This is especially important as metabarcoding coupled with deep sequencing and specific primer sets has shown to be more sensitive than single species qPCR detections, providing a superior tool for biodiversity analysis of rare targets^39,40^.

## MATERIALS AND METHODS

### Fish sampling

British Columbia (BC) has a total of approximately 92 (75 native and 17 invasive) species of fish that use freshwater for all or part of their life cycle, including significant geographical variants or subspecies^41–44^. We collected samples from at least one individual for 85 of these, either from tissue samples (frozen or in ethanol) and/or DNA from curated voucher specimens. As priority, voucher specimens were collected from within their BC range (n=35), and secondarily from other areas of Canada or the USA (Supplementary Table 1). This included Washington (n=4), Alaska (n=2), Oregon (n=1), Yukon (n=9), Alberta (n=3), Manitoba (n=1), Ontario (n=17), and Quebec (n=12). Details on voucher specimens are in the quality control section below. For species with evolutionary significant geographic variants or subspecies, we collected one individual from each distinct taxon. These include westslope cutthroat trout (*Oncorhynchus clarkii ssp. lewisi*)^42^, Coastal and Interior lineages of bull trout (*Salvelinus confluentus*)^44^, and Morrison Creek Lamprey (*Lampetra richardsoni var. marifuga*)^41^.

Shotgun DNA sequencing, mitogenome and rDNA assemblies, and annotation Total genomic DNA was extracted from fin or muscle tissue using the Qiagen DNeasy Blood and Tissue kit and quantified using Quant-iT™ PicoGreen Assay (ThermoFisher). Input amounts normalized to 10 ng were used to build Illumina DNA libraries, which were sequenced on a NovaSeq SP flow cell (2 × 250bp) at a target sequencing depth of 5 million reads per sample. For the subset of samples with mitochondrial genome and nuclear rDNA cistron coverage below 20-fold after the first run, the same library was sequenced using another Illumina NovaSeq SP flow cell (2 × 250 bp kit) to increase read depth by 1 to 14M reads per sample. For the subset of samples for which mitochondrial genome assembly was not possible after the first run, the original genomic DNA was used in a secondary independent Illumina DNA library preparation with minor modifications for low DNA input. This was then sequenced using the Illumina NovaSeq SP flow cell (2 × 250 bp kit) at a target sequencing depth of 10 million reads per sample.

Raw sequencing data were demultiplexed and trimmed of indices using Illumina’s bcl2fastq (version 2.20.0.422) software. For each sample, trimmomatic (version 0.39)^45^ was used to remove Illumina adapters and trim low-quality base calls from the ends of reads. We used several de novo assembly/annotation software toolkits since no single tool was successful at analyzing all samples, as detailed here. GetOrganelle (version 1.7.4.1)^46^ was used to assemble paired-end reads into whole mitogenomes (or scaffolds when complete assembly was not possible); then MitoZ (version 2.3)^47^ was used to annotate assemblies or scaffolds. Alternatively, some samples were assembled and annotated using MitoFlex (version 0.2.9)^48^, and others were assembled using the de novo assembler ABySS (version 2.2.5)^49^ and then annotated using MitoZ. Nuclear rDNA regions were annotated using the software barrnap (version 0.9)^50^. We included all results where the length was at least 95% complete.

### Quality assurance

To ensure quality and defensibility of information in the NAMERS database, we aimed to satisfy three criteria for all samples sequenced: (1) availability of a museum-catalogued voucher specimen; (2) minimum voucher specimen metadata consisting of collection site name, geographic location, sampling date, and the name and affiliation of who did the morphological identification; and (3) genetic species identity verification using COI barcode sequences. To ensure traceability of sequence data to physical voucher specimens and minimize the likelihood of sequencing misidentified material, tissues to be sequenced were predominantly sourced from museum collections, as follows: the Royal Ontario Museum (n=45); the University of British Columbia’s Beaty Biodiversity Museum (n=13); and the University of Washington Burke Museum Ichthyology Collection (n=5). Exceptions to this included 9 tissue samples from the Beaty Biodiversity Museum, collected and identified by fish collection director Dr. Eric B. Taylor (E. Taylor, pers. comm.); six of which have voucher specimens that are not catalogued and four of which had no voucher specimen (both bull trout lineages, *Salvelinus confluentus*; lake trout, *Salvelinus namaycush*; and longnose sucker, *Catostomus catostomus*). Tissue from the inconnu (*Stenodus leucichthys*), collected by the Teslin Tlingit Council, also does not have a whole voucher specimen but has a tissue voucher housed at the Pacific Biological Station (Nanaimo, BC). The remainder of voucher specimens were collected by Fisheries and Oceans Canada (n=19) and are currently being catalogued at the Royal British Columbia Museum. All voucher information is included on the species pages for each specimen in NAMERS and in Supplementary Table 1.

To verify concordance between morphological taxonomy and molecular taxonomy, whole COI sequences from each mitogenome were manually aligned and inspected with DNA barcodes produced by Hubert et al.^51^ in the Canadian Freshwater Fish Barcode Database (BCF and BCFB projects in BOLD, n= 190 species). We used a divergence cut-off of 0.6% to confirm matches, as Hubert et al.^51^ calculated a maximum intraspecific genetic distance of 0.6% in the ~650 bp COI barcode region for Canadian freshwater fish (excluding outliers; note that average interspecific divergence was a much higher 7.5%^51^).

The exception was lampreys (Petromyzontidae), which are not in the Canadian Freshwater Fish Barcode Database^51^ and have very little interspecific mitogenomic variation. Thus we took a multi-faceted approach to verify the genetic identity of the five species in NAMERS, including first verifying that each NAMERS sequence matched a record in GenBank derived from a voucher specimen. The *Entosphenus* genus contains *E. tridentatus* and *E. macrostomus*, the latter of which is found only in Cowichan Lake, hence we use sampling location to identify this specimen. *E. tridentatus* is widespread and the NAMERS specimen was collected in marine waters and identified by presence of three prominent teeth by lamprey expert Joy Wade (Joy Wade, pers., comm.). The *Lampetra* genus contains geographically isolated *L. richardsoni var. marifuga*, for which the NAMERS specimen was collected within its range at Morrison Creek, BC, Canada. It also contains *L. ayresii* and *L. richardsoni* which were differentiated based on their non-overlapping habitat use: the former was collected as an adult in marine waters whereas the latter in freshwater.

### A mitogenome portal for applied eDNA metabarcoding

NAMERS sequences and associated metadata were deposited in GenBank (under the BC Freshwater Fish Genome Project) and in a newly developed, purpose-built online mitogenome and nuclear rDNA cistron sequence data portal specifically for applied eDNA metabarcoding (https://namers.ca). Specific functionalities of the portal are summarized in Results. Assessments of amplification and taxonomic resolution efficiencies of genetic markers are critical for sound applied eDNA metbarcoding study design as both are key determinants of success^14,23^. To illustrate the value of mitogenome data for marker selection, we used NAMERS to evaluate the efficacy of existing mitochondrial fish metabarcoding markers for both amplifying targets and providing species level resolution. We selected 19 eDNA metabarcoding markers targeting teleost fish from across four mitochondrial gene regions (12S, n=7; 16S, n=5; COI, n=4; and cytochrome b (CYTB), n=3) from the literature, all of which amplify targets smaller than the maximum allowable length for Illumina sequencing kits (Table 1). Each primer pair was aligned and manually checked for the number of mismatches with each species; degenerate bases in primers were scored as mismatches if none of the nucleotides coded for by the degeneracy matched the species nucleotide at that site. Amplicons for each species were generated and checked for uniqueness using two methods in R^52^. First, the haplotype function from the haplotypes package^53^ identified unique haplotypes based on genetic distance calculated using dist.dna with model “raw” and pairwise deletion of indels in the ape package^54^. This method did not identify unique haplotypes due to indels, therefore, the GetHaplo function in the SIDIER package^55^ was used to identify these. Each unique haplotype was manually checked and considered as providing species level resolution if it was at least 1 bp different (including indels) from another species.

## Acknowledgements

We would like to thank the following people for contributions to the study with specimen collection and curation. From DFO; Liane Stenhouse, Paul Grant, Nellie Gagné, Louis Marie Roux, Mélanie Roy. Joy Wade (Fundy Aquaculture Services), Rick Taylor (UBC Beatty Museum of Biodiversity), Jordan Rosenfeld (BC Ministry of Environment), Bob Hanner (University of Guelph), Daniel Heath (University of Windsor), Royal Ontario Museum, Gavin Hanke (Royal BC Museum), Teslin Tlingit Council, Pascale Savage (Yukon Government), Caren Helbing (University of Victoria), Amelia Louden (Burke Museum), and Louis Lopez (University of Victoria). This research was funded by Genome BC, Project #SIP26-06.

## Author Contributions

KMW, CLA, and SRG conceived of the study and obtained project funding. KMW prepared samples for sequencing. MH, NF, and GACS managed sequencing and performed bioinformatics. MK and GACS built the website with input from all authors. KMW and CLA wrote the manuscript with input from all authors.

## Data Availability

All processed genetic data is available from https://namers.ca and is available in Genbank (Accession Numbers in Supplementary Tables 2 and 3).

## Additional Information

The authors declare no competing interests.

## LITERATURE CITED

1. Cordier, T. et al.. Ecosystems monitoring powered by environmental genomics: A review of current strategies with an implementation roadmap. Mol. Ecol. 30, 2937–2958 (2021).

2. Cristescu, M. E. & Hebert, P. D. N. Uses and Misuses of Environmental DNA in Biodiversity Science and Conservation. Annu. Rev. Ecol. Evol. Syst. 49, 209–230 (2018).

3. Hering, D. et al.. Implementation options for DNA-based identification into ecological status assessment under the European Water Framework Directive. Water Res. 138, 192–205 (2018).

4. Ruppert, K. M., Kline, R. J. & Rahman, M. S. Past, present, and future perspectives of environmental DNA (eDNA) metabarcoding: A systematic review in methods, monitoring, and applications of global eDNA. Glob. Ecol. Conserv. 17, e00547 (2019).

5. Schenekar, T. The current state of eDNA research in freshwater ecosystems: are we shifting from the developmental phase to standard application in biomonitoring? Hydrobiologia (2022) doi:10.1007/s10750-022-04891-z.

6. Wang, S. et al.. Methodology of fish eDNA and its applications in ecology and environment. Sci. Total Environ. 755, (2021).

7. Zaiko, A., Pochon, X., Garcia-Vazquez, E., Olenin, S. & Wood, S. A. Advantages and limitations of environmental DNA/RNA tools for marine biosecurity: Management and surveillance of non-indigenous species. Front. Mar. Sci. 5, (2018).

8. Baird, D. J. & Hajibabaei, M. Biomonitoring 2.0: A new paradigm in ecosystem assessment made possible by next-generation DNA sequencing. Mol. Ecol. 21, 2039–2044 (2012).

9. Zinger, L. et al.. DNA metabarcoding—Need for robust experimental designs to draw sound ecological conclusions. Mol. Ecol. 28, 1857–1862 (2019).

10. Darling, J. A. & Mahon, A. R. From molecules to management: Adopting DNA-based methods for monitoring biological invasions in aquatic environments. Environ. Res. 111, 978–988 (2011).

11. Darling, J. A. How to learn to stop worrying and love environmental DNA monitoring. Aquat. Ecosyst. Health Manag. 22, 440–451 (2019).

12. Darling, J. A., Pochon, X., Abbott, C. L., Inglis, G. J. & Zaiko, A. The risks of using molecular biodiversity data for incidental detection of species of concern. Divers. Distrib. 26, 1116–1121 (2020).

13. Hajibabaei, M., Shokralla, S., Zhou, X., Singer, G. A. C. & Baird, D. J. Environmental Barcoding: A Next-Generation Sequencing Approach for Biomonitoring Applications Using River Benthos. PLoS One 6, e17497 (2011).

14. Taberlet, P., Coissac, E., Pompanon, F., Brochmann, C. & Willerslev, E. Towards nextgeneration biodiversity assessment using DNA metabarcoding. Mol. Ecol. 21, 2045–2050 (2012).

15. Ratnasingham, S. & Hebert, P. D. N. BOLD: The Barcode of Life Data System: Barcoding. Mol. Ecol. Notes 7, 355–364 (2007).

16. Benson, D. A. et al.. GenBank. Nucleic Acids Res. 28, 15–18 (2000).

17. Dziedzic, E. et al.. Transitioning from environmental genetics to genomics using mitogenome reference databases. (2022) doi:10.22541/au.164873596.68312121/v1.

18. Schroeter, J. C., Maloy, A. P., Rees, C. B. & Bartron, M. L. Fish mitochondrial genome sequencing: expanding genetic resources to support species detection and biodiversity monitoring using environmental DNA. Conserv. Genet. Resour. 12, 433–446 (2020).

19. Leese, F. et al. Chapter Two - Why We Need Sustainable Networks Bridging Countries, Disciplines, Cultures and Generations for Aquatic Biomonitoring 2.0: A Perspective Derived From the DNAqua-Net COST Action. in Next Generation Biomonitoring: Part 1 (eds. Bohan, D. A., Dumbrell, A. J., Woodward, G. & Jackson, M. B. T.-A. in E. R.) vol. 58 63–99 (Academic Press, 2018).

20. Mathieu, C. et al.. A Systematic Review of Sources of Variability and Uncertainty in eDNA Data for Environmental Monitoring. Frontiers in Ecology and Evolution vol. 8 at https://www.frontiersin.org/articles/10.3389/fevo.2020.00135 (2020).

21. Ficetola, G. F. et al.. Comparison of markers for the monitoring of freshwater benthic biodiversity through DNA metabarcoding. Mol. Ecol. 30, 3189–3202 (2021).

22. Piñol, J., Mir, G., Gomez-Polo, P. & Agustí, N. Universal and blocking primer mismatches limit the use of high-throughput DNA sequencing for the quantitative metabarcoding of arthropods. Mol. Ecol. Resour. 15, 819–830 (2015).

23. Deagle, B. E., Jarman, S. N., Coissac, E., Pompanon, F. & Taberlet, P. DNA metabarcoding and the cytochrome c oxidase subunit I marker: not a perfect match. Biol. Lett. 10, (2014).

24. Creer, S. et al.. The ecologist’s field guide to sequence-based identification of biodiversity. Methods Ecol. Evol. 7, 1008–1018 (2016).

25. McElroy, M. E. et al.. Calibrating Environmental DNA Metabarcoding to Conventional Surveys for Measuring Fish Species Richness. Front. Ecol. Evol. 8, (2020).

26. Collins, R. A. et al.. Non-specific amplification compromises environmental DNA metabarcoding with COI. Methods Ecol. Evol. 10, 1985–2001 (2019).

27. Edgar, R. C. MUSCLE: multiple sequence alignment with high accuracy and high throughput. Nucleic Acids Res. 32, 1792–1797 (2004).

28. Vences, M. et al.. Freshwater vertebrate metabarcoding on Illumina platforms using double-indexed primers of the mitochondrial 16S rRNA gene. Conserv. Genet. Resour. 8, 323–327 (2016).

29. Mauvisseau, Q. et al.. Influence of accuracy, repeatability and detection probability in the reliability of species-specific eDNA based approaches. Sci. Rep. 9, 580 (2019).

30. Darling, J. A., Jerde, C. L. & Sepulveda, A. J. What do you mean by false positive? Environ. DNA 3, 879–883 (2021).

31. Zaiko, A. et al.. Towards reproducible metabarcoding data: Lessons from an international cross-laboratory experiment. Mol. Ecol. Resour. 22, 519–538 (2022).

32. Collins, R. A. et al.. Meta-Fish-Lib: A generalised, dynamic DNA reference library pipeline for metabarcoding of fishes. J. Fish Biol. 99, 1446–1454 (2021).

33. Weigand, H. et al.. DNA barcode reference libraries for the monitoring of aquatic biota in Europe: Gap-analysis and recommendations for future work. Sci. Total Environ. 678, 499–524 (2019).

34. Froese, R. & Pauly, D. FishBase. FishBase 2000: concepts, design and data sources 1–344 (2000).

35. Gold, Z. et al.. Improving metabarcoding taxonomic assignment: A case study of fishes in a large marine ecosystem. Mol. Ecol. Resour. 21, 2546–2564 (2021).

36. Feist, S. M. & Lance, R. F. Advanced molecular-based surveillance of quagga and zebra mussels: A review of environmental DNA/RNA (eDNA/eRNA) studies and considerations for future directions. NeoBiota 66, 117–159 (13AD).

37. Hoban, M. L. et al.. Skimming for barcodes: rapid production of mitochondrial genome and nuclear ribosomal repeat reference markers through shallow shotgun sequencing. PeerJ 10, e13790 (2022).

38. Hajibabaei, M. Demystifying eDNA validation. Trends Ecol. Evol. 37, 826–828 (2022).

39. McCarthy, A. et al.. Comparative analysis of fish environmental DNA reveals higher sensitivity achieved through targeted sequence-based metabarcoding. Mol. Ecol. Resour. n/a, (2022).

40. Westfall, K. M., Therriault, T. W. & Abbott, C. L. Targeted next-generation sequencing of environmental DNA improves detection of invasive European green crab (Carcinus maenas). Environ. DNA 4, 440–452 (2022).

41. Beamish, R. J. Evidence that parasitic and non-parasitic life history types are produced by one population of lamprey. Can. J. Fish. Aquat. Sci. 44, 1779–1782 (1987).

42. McPhail, J. D. The Freshwater Fishes of British Columbia. vol. 121 (2007).

43. Ruskey, J. A. & Taylor, E. B. Morphological and genetic analysis of sympatric dace within the Rhinichthys cataractae species complex: a case of isolation lost. Biol. J. Linn. Soc. 117, 547–563 (2016).

44. Taylor, E. B., Pollard, S. & Louie, D. Mitochondrial DNA variation in bull trout (Salvelinus confluentus) from northwestern North America: implications for zoogeography and conservation. Mol. Ecol. 8, 1155–1170 (1999).

45. Bolger, A. M., Lohse, M. & Usadel, B. Trimmomatic: a flexible trimmer for Illumina sequence data. Bioinformatics 30, 2114–2120 (2014).

46. Jin, J.-J. et al.. GetOrganelle: a fast and versatile toolkit for accurate de novo assembly of organelle genomes. Genome Biol. 21, 241 (2020).

47. Meng, G., Li, Y., Yang, C. & Liu, S. MitoZ: A toolkit for animal mitochondrial genome assembly, annotation and visualization. Nucleic Acids Res. 47, (2019).

48. Li, J.-Y., Li, W.-X., Wang, A.-T. & Zhang, Y. MitoFlex: an efficient, high-performance toolkit for animal mitogenome assembly, annotation and visualization. Bioinformatics 37, 3001–3003 (2021).

49. Simpson, J. T. et al.. ABySS: A parallel assembler for short read sequence data. Genome Res. 19, 1117–1123 (2009).

50. Seeman, T. barnnap. at https://github.com/tseemann/barrnap (2009).

51. Hubert, N. et al.. Identifying Canadian Freshwater Fishes through DNA Barcodes. PLoS One 3, e2490 (2008).

52. R Core Team, R. R: A Language and Environment for Statistical Computing. at (2018).

53. Atkas, C. haplotypes: Manipulating DNA Sequences and Estimating Unambiguous Haplotype Network with Statistical Parsimony. at https://cran.r-project.org/web/packages/haplotypes/index.html (2020).

54. Paradis, E. & Schliep, K. ape 5.0: an environment for modern phylogenetics and evolutionary analyses in R. Bioinformatics bty633–bty633 (2018).

55. Muñoz-Pajares, A. J. Nosidier: Substitution and Indel Distances to Infer Evolutionary Relationships. at https://rdrr.io/cran/sidier/ (2021).

