## Supplementary Table 1 for "NAMERS: a purpose-built reference DNA sequence database to support applied eDNA metabarcoding"

| **Family** | **Genus** | **Species** | **Specimen_ID** | **Country** | **PROVINCE/STATE** | **Site** | **Year** | **Institution** | **Catalogue Number** |
| --- | --- | --- | --- | --- | --- | --- | --- | --- | --- |
| Acipenseridae | Acipenser | medirostris | AcMed | USA | OR | Gray's Harbour | 2021 | WDFW | 12NN0243 |
| Acipenseridae | Acipenser | transmontanus | AcTra | Canada | BC | Vancouver Island University (holding facility) | 2013 | ROM | T14631-94838 |
| Catostomidae | Catostomus | catostomus | CaCat | Canada | BC | Red Rock Creek | 2000 | UBC | X |
| Catostomidae | Catostomus | commersonii | CaCom | Canada | QC | Riviere Sauvage tributary | 2005 | ROM | T01087-80075 |
| Catostomidae | Catostomus | macrocheilus | CaMac | Canada | BC | West Kettle River | 2020 | PBS | JW78/JWDNA41 |
| Catostomidae | Pantosteus | bondi | PaBon | Canada | BC | Similkameen River | N/A | UBC | No Catalogue |
| Catostomidae | Pantosteus | jordani | PaJon | Canada | AB | Unknown | 2012 | ROM | T13372-93986 |
| Centrarchidae | Lepomis | gibbosus | LeGib | Canada | ON | Lake Erie | 2005 | ROM | T00793-79869 |
| Centrarchidae | Lepomis | macrochirus | LeMac | Canada | ON | Opinicon Lake | 2005 | ROM | T01069-80072 |
| Centrarchidae | Micropterus | dolomieu | MiDol | Canada | QC | Lac Saint-Louis | 2005 | ROM | T01187-80102 |
| Centrarchidae | Micropterus | salmoides | MiSal | Canada | ON | Lac Saint-Louis | 2005 | ROM | T01066-80071 |
| Centrarchidae | Pomoxis | nigromaculatus | PoNir | Canada | ON | Lake Simcoe | 1994 | ROM | T00483-75658 |
| Clupeidae | Alosa | sapidissima | AlSap | Canada | QC | Lac Saint-Pierre | 2005 | ROM | T01250-80125 |
| Cobittidae | Misgurnus | anguillicaudatus | MiAng | Canada | N/A | Petculture Dieppe | 2015 | ROM | T07232-101514 |
| Cottidae | Cottus | aleuticus | CoAle | USA | WA | Mason (Coulter creek tributary) | 2009 | UW | UW 151939 |
| Cottidae | Cottus | asper | CoAsp | Canada | BC | Little River | 2020 | PBS | JW174/JWDNA92 |
| Cottidae | Cottus | cognatus | CoCog | Canada | YT | Porcupine River tributary | 2005 | ROM | T02204-78455 |
| Cottidae | Cottus | confuses | CoCon | USA | WA | Twin Creek | 2011 | UW | UW 152071 |
| Cottidae | Cottus | hubbsi | CoHub | Canada | BC | Summer's creek | 2011 | UBC | BC 11-0641 |
| Cottidae | Cottus | rhotheus | CoRho | Canada | BC | Little Slocan River | 2000 | UBC | BC 11-0036 |
| Cottidae | Cottus | ricei | CoRic | Canada | QC | St. Lawrence River | 2006 | ROM | T03576-81216 |
| Cottidae | Leptocottus | armatus | LeArm | Canada | BC | Whiffin Spot | 2019 | PBS | WS12 |
| Cyprinidae | Acrocheilus | alutaceus | AcAlu | Canada | BC | Kettle River | 2020 | PBS | JW87/JWDNA44 |
| Cyprinidae | Carassius | auratus | CaAur | Canada | ON | Dundas Pond | 2013 | ROM | T13358-93983 |
| Cyprinidae | Chrosomus | eos | ChEos | Canada | ON | Sauble River | 2007 | ROM | T10980-89468 |
| Cyprinidae | Chrosomus | neogaeus | ChNeo | Canada | ON | Sauble River | 2007 | ROM | T10976-89466 |
| Cyprinidae | Couesius | plumbeus | CoPlu | Canada | ON | Lake Huron | 2005 | ROM | T00531-79889 |
| Cyprinidae | Cyprinus | carpio | CyCar | Canada | ON | Big Creek | 1997 | ROM | T05535-70937 |
| Cyprinidae | Hybognathus | hankinsoni | HyHan | Canada | BC | Bog pond | 2005 | ROM | T04561 |
| Cyprinidae | Margariscus | nachtriebi | MaNac | Canada | BC | McConchie Creek | 2008 | UBC | No Catalogue |
| Cyprinidae | Notropis | atherinoides | NoAth | Canada | QC | Lac Saint-Louis | 2005 | ROM | T01472-80191 |
| Cyprinidae | Notropis | hudsonius | NoHud | Canada | ON | Lake Erie | 1997 | ROM | T05662-70972 |
| Cyprinidae | Pimephales | promelas | PiPro | Canada | ON | Humber River | 1993 | ROM | T05845 |
| Cyprinidae | Platygobio | gracilis | PlGra | Canada | AB | Unknown | 2012 | ROM | T13370-95503 |
| Cyprinidae | Ptychocheilus | oregonensis | PtOre | Canada | BC | Kettle River | 2020 | PBS | JW35/JWDNA20 |
| Cyprinidae | Rhinichthys | cataractae | RhCat | Canada | ON | Penetangore River | 2007 | ROM | T10943-89441 |
| Cyprinidae | Rhinichthys | osculus | RhOsc | Canada | BC | Kettle River | 2020 | PBS | JW11/JWDNA6 |
| Cyprinidae | Rhinichthys | falcatus | RhFal | Canada | BC | Similkameen River | 2011 | UBC | BC 12-039 |
| Cyprinidae | Richardsonius | balteatus | RiBal | Canada | BC | West Kettle River | 2020 | PBS | JW130/JWDNA58 |
| Cyprinidae | Tinca | tinca | TiTin | Canada | BC | Kootenay or Okanagan Region | 2017 | UBC | No Catalogue |
| Embiotocidae | Cymatogaster | aggregata | CyAgg | Canada | BC | Maple Bay | 2019 | PBS | MB-s01-1 |
| Esocidae | Esox | lucius | EsLuc | Canada | QC | Lac Saint-Paul | 2005 | ROM | T01147-80092 |
| Gasterosteidae | Culaea | inconstans | CuInc | Canada | QC | Riviere Saint-Jean | 2005 | ROM | T01126-80086 |
| Gasterosteidae | Gasterosteus | aculeatus | GaAcu | Canada | BC | Whiffin Spot | 2019 | PBS | WS05 |
| Gasterosteidae | Pungitius | pungitius | PuPun | Canada | QC | St Lawrence River | 2005 | ROM | T01850-80265 |
| Hiodontidae | Hiodon | alosoides | HiAlo | Canada | MB | Lake Winnipeg | 2012 | ROM | T13388-93987 |
| Ictaluridae | Ameiurus | melas | AmMel | Canada | BC | Osoyoos Lake | 2005 | ROM | T02262-81212 |
| Ictaluridae | Ameiurus | natalis | AmNat | Canada | ON | Tumblesons Pond | 1998 | ROM | T03628-71710 |
| Ictaluridae | Ameiurus | nebulosus | AmNeb | Canada | QC | Lac Saint-Pierre | 2005 | ROM | T01231-80118 |
| Lotidae | Lota | lota | LoLot | Canada | BC | Teslin Lake | 2005 | ROM | T02264-81208 |
| Osmeridae | Osmerus | mordax | OsMor | Canada | ON | Lake Erie | 2000 | ROM | T00005-75473 |
| Osmeridae | Spirinchus | thaleichthys | SpTha | Canada | BC | Pitt Lake | 2006 | UBC | No Catalogue |
| Osmeridae | Thaleichthys | pacificus | ThPac | Canada | BC | DFO-2018-18 set 7 | 2018 | PBS | JW176/JWDNA94 |
| Percidae | Perca | flavescens | PeFla | Canada | QC | Riviere Laval | 2005 | ROM | T01193-80105 |
| Percidae | Sander | vitreus | SaVit | Canada | QC | Lac Saint-Louis | 2005 | ROM | T01224-80117 |
| Percopsidae | Percopsis | omiscomaycus | PeOmi | Canada | ON | Thames River | 2005 | ROM | T00548-79871 |
| Petromyzontidae | Entosphenus | macrostomus | EnMac | Canada | BC | Cowichan Lake | 2013 | ROM | T14630-94828 |
| Petromyzontidae | Entosphenus | tridentatus | EnTri | Canada | BC | Franklin 2020-02 | 2020 | PBS | JW191/JWDNA109 |
| Petromyzontidae | Lampetra | ayresii | LaAyr | Canada | BC | Franklin 2020-02 | 2020 | PBS | JW186/JWDNA104 |
| Petromyzontidae | Lampetra | richardsoni | LaRic | USA | WA | Black jack creek (Kitsap) | 2009 | UW | UW 152225 |
| Petromyzontidae | Lampetra | richardsoni var. marifuga | LaAyrMC | Canada | BC | Morrison Creek | 2020 | PBS | JW175/JWDNA93 |
| Pleuronectidae | Platichthys | stellatus | PlSte | Canada | BC | Murder Bay | 2019 | PBS | MUB10-1 |
| Salmonidae | Coregonus | artedi | CoArt | Canada | ON | Lake Erie | 1999 | ROM | T03648-72207 |
| Salmonidae | Coregonus | clupeaformis | CoClu | Canada | YT | Porcupine River tributary | 2005 | ROM | T02206-78456 |
| Salmonidae | Coregonus | nasus | CoNas | Canada | YT | Dawson City | 2005 | UBC | No Catalogue |
| Salmonidae | Coregonus | sardinella | CoSar | Canada | YT | Dawson City | 2005 | UBC | No Catalogue |
| Salmonidae | Oncorhynchus | clarki | OnCla | Canada | BC | Morrison Creek | 2020 | PBS | JW160/JWDNA78 |
| Salmonidae | Oncorhynchus | clarkii ssp. lewisi | OnClaL | Canada | AB | Girardi Creek | 2017 | Uvic | ONCLle-v1 |
| Salmonidae | Oncorhynchus | gorbuscha | OnGor | USA | AK | Gulf of Alaska | 2011 | UW | UW 151072 |
| Salmonidae | Oncorhynchus | keta | OnKet | Canada | BC | Sidney Inlet | 2013 | ROM | T14659-94840 |
| Salmonidae | Oncorhynchus | kisutch | OnKis | Canada | ON | Wilmot Creek | 1994 | ROM | T03344-68277 |
| Salmonidae | Oncorhynchus | mykiss | OnMyk | Canada | YT | McIntrye Creek | 1994 | ROM | T02789-68289 |
| Salmonidae | Oncorhynchus | nerka | OnNer | Canada | BC | Franklin no info | 2020 | PBS | JW192/JWDNA110 |
| Salmonidae | Oncorhynchus | tshawytscha | OnTsh | Canada | YT | Klondike River | 2006 | ROM | T02243-79128 |
| Salmonidae | Prosopium | coulterii | PrCou | USA | AK | Chignik Lake | 2008 | PBS | 2353-005 (tissue vial #) |
| Salmonidae | Prosopium | cylindraceum | PrCyl | Canada | YT | Ensley Creek | 2006 | ROM | T03792 |
| Salmonidae | Prosopium | williamsoni | PrWil | USA | WA | Bacon Creek | 1993 | UW | UW 048051 |
| Salmonidae | Salmo | salar | SaSal | Canada | BC | Koksilah | 2021 | PBS | GBC 30 |
| Salmonidae | Salvelinus | confluentus (Coastal lineage) | SaConC | Canada | BC | Pitt Lake | 1999 | UBC | X |
| Salmonidae | Salvelinus | confluentus (Interior lineage) | SaConI | Canada | BC | Dease Lake | 1999 | UBC | X |
| Salmonidae | Salvelinus | fontinalis | SaFon | Canada | QC | Riviere Trinite | 2005 | ROM | T01542-80198 |
| Salmonidae | Salvelinus | malma | SaMal | Canada | BC | Morrison Creek | 2020 | PBS | JW 165/JWDNA 83 |
| Salmonidae | Salvelinus | namaycush | SaNam | Canada | BC | Nation Lakes | 1998 | UBC | X |
| Salmonidae | Stenodus | leucichthys | StLeu | Canada | YT | Teslin Lake | 2021 | PBS | EDI2021DNA064 |
| Salmonidae | Thymallus | arcticus | ThArc | Canada | YT | Burwash Creek | 2006 | ROM | T02236-79136 |
| ROM = Royal Ontario Museum | |  |  |  |  |  |  |  |  |
| UBC = University of British Columbia Beatty Museum of Biodiversity | | |  |  |  |  |  |  |  |
| UW = University of Washington Fish Collection | |  |  |  |  |  |  |  |  |
| Uvic = University of Victoria, Prof. Caren Helbing collection | | |  |  |  |  |  |  |  |
| PBS = Pacific Biological Station, Fisheries and Oceans Canada | | |  |  |  |  |  |  |  |
| WDFW = Washington Department of Fish and Wildlife Fish Collection | | |  |  |  |  |  |  |  |
| No Catalogue = voucher specimen permanently housed in organismal or molecular collection at Beaty Museum of Biodiversity (UBC) but not catalogued | | | | | | |  |  |  |
| X = no voucher specimen but verified visually and through molecular diagnostics | | |  |  |  |  |  |  |  |
