## Supplementary Table 2 for "NAMERS: a purpose-built reference DNA sequence database to support applied eDNA metabarcoding"

| **Family** | **Genus** | **Species** | **Mt genome length (kbp)** | **Nuclear rDNA** |  |  | **Complete** |  |  |  |
| --- | --- | --- | --- | --- | --- | --- | --- | --- | --- | --- |
|  |  |  |  | **5.8S** | **18S** | **28S** | **Mitogenome** |  |  |  |
| Acipenseridae | *Acipenser* | *medirostris* | 16.690 | OR020751 | OR051928 | OR020610 | OR637236 |  |  |  |
| Acipenseridae | *Acipenser* | *transmontanus* | 16.529 |  | OR051929 | OR020611 | OR637237 |  |  |  |
| Catostomidae | *Catostomus* | *catostomus* | 16.580 | OR020752 | OR051930 | OR020612 | OR637238 |  | LEGEND |  |
| Catostomidae | *Catostomus* | *commersonii* | 16.622 | OR020753 | OR051931 | OR020613 | OR637239 |  |  | full |
| Catostomidae | *Catostomus* | *macrocheilus* | 16.626 | OR020754 | OR051932 | OR020614 | OR637240 |  | XX123456 | partial |
| Catostomidae | *Pantosteus* | *bondi* | N/A | OR020755 | OR051933 | OR020615 | N/A |  |  | none |
| Catostomidae | *Pantosteus* | *jordani* | 16.623 | OR020756 | OR051934 | OR020616 | OR637241 |  |  |  |
| Centrarchidae | *Lepomis* | *gibbosus* | 16.504 | OR020757 | OR051935 | OR020617 | OR637242 |  |  |  |
| Centrarchidae | *Lepomis* | *macrochirus* | 16.491 | OR020758 | OR051936 | OR020618 | OR637243 |  |  |  |
| Centrarchidae | *Micropterus* | *dolomieu* | 16.486 | OR020759 | OR051937 | OR020619 | OR637244 |  |  |  |
| Centrarchidae | *Micropterus* | *salmoides* | 16.489 | OR020760 | OR051938 | OR020620 | OR637245 |  |  |  |
| Centrarchidae | *Pomoxis* | *nigromaculatus* | 16.497 | OR020761 | OR051939 | OR020621 | OR637246 |  |  |  |
| Clupeidae | *Alosa* | *sapidissima* | 18.142 | OR020762 | OR051940 | OR020622 | OR637247 |  |  |  |
| Cobittidae | *Misgurnus* | *anguillicaudatus* | 16.647 | OR020763 | OR051941 | OR020623 | OR637248 |  |  |  |
| Cottidae | *Cottus* | *aleuticus* | 16.603 | OR020764 | OR051942 | OR020624 | OR637249 |  |  |  |
| Cottidae | *Cottus* | *asper* | 16.821 | OR020765 | OR051943 | OR020625 | OR637250 |  |  |  |
| Cottidae | *Cottus* | *cognatus* | 16.539 | OR020766 | OR051944 | OR020626 | OR637251 |  |  |  |
| Cottidae | *Cottus* | *confuses* | 16.532 | OR020767 | OR051945 | OR020627 | OR637252 |  |  |  |
| Cottidae | *Cottus* | *hubbsi* | 16.534 | OR020768 | OR051946 | OR020628 | OR637253 |  |  |  |
| Cottidae | *Cottus* | *rhotheus* | 16.739 | OR020769 | OR051947 | OR020629 | OR637254 |  |  |  |
| Cottidae | *Cottus* | *ricei* | 16.533 | OR020770 | OR051948 | OR020630 | OR637255 |  |  |  |
| Cottidae | *Leptocottus* | *armatus* | 16.565 | OR020771 | OR051949 | OR020631 | OR637256 |  |  |  |
| Cyprinidae | *Acrocheilus* | *alutaceus* | 16.592 | OR020772 | OR051950 | OR020632 | OR637257 |  |  |  |
| Cyprinidae | *Carassius* | *auratus* | 16.513 | OR020773 | OR051951 |  | OR637258 |  |  |  |
| Cyprinidae | *Chrosomus* | *eos* | 16.624 | OR020774 | OR051952 | OR020633 | OR637259 |  |  |  |
| Cyprinidae | *Chrosomus* | *neogaeus* | 16.607 | OR020775 | OR051953 | OR020634 | OR637260 |  |  |  |
| Cyprinidae | *Couesius* | *plumbeus* | 16.605 | OR020776 | OR051954 | OR020635 | OR637261 |  |  |  |
| Cyprinidae | *Cyprinus* | *carpio* | 16.531 | OR020777 | OR051955 | OR020636 | OR637262 |  |  |  |
| Cyprinidae | *Hybognathus* | *hankinsoni* | 16.699 | OR020778 | OR051956 | OR020637 | OR637263 |  |  |  |
| Cyprinidae | *Margariscus* | *nachtriebi* | N/A |  | OR051957 |  | N/A |  |  |  |
| Cyprinidae | *Notropis* | *atherinoides* | 16.744 | OR020779 | OR051958 |  | OR637264 |  |  |  |
| Cyprinidae | *Notropis* | *hudsonius* | 16.659 | OR020780 |  | OR020638 | OR637265 |  |  |  |
| Cyprinidae | *Pimephales* | *promelas* | 16.721 | OR020781 | OR051959 | OR020639 | OR637266 |  |  |  |
| Cyprinidae | *Platygobio* | *gracilis* | 16.985 | OR020782 | OR051960 | OR020640 | OR637267 |  |  |  |
| Cyprinidae | *Ptychocheilus* | *oregonensis* | 16.604 | OR020783 | OR051961 | OR020641 | OR637268 |  |  |  |
| Cyprinidae | *Rhinichthys* | *cataractae* | 16.660 | OR020784 | OR051962 | OR020642 | OR637269 |  |  |  |
| Cyprinidae | *Rhinichthys* | *osculus* | 16.655 | OR020785 | OR051963 | OR020643 | OR637270 |  |  |  |
| Cyprinidae | *Rhinichthys* | *falcatus* | 16.766 | OR020786 | OR051964 | OR020644 | OR637271 |  |  |  |
| Cyprinidae | *Richardsonius* | *balteatus* | 16.647 | OR020787 | OR051965 | OR020645 | OR637272 |  |  |  |
| Cyprinidae | *Tinca* | *tinca* | 16.614 | OR020788 |  | OR020646 | OR637273 |  |  |  |
| Embiotocidae | *Cymatogaster* | *aggregata* | 17.656 | OR020789 | OR051966 | OR020647 | OR637274 |  |  |  |
| Esocidae | *Esox* | *lucius* | 16.125 | OR020790 | OR051967 | OR020648 | OR637275 |  |  |  |
| Gasterosteidae | *Culaea* | *inconstans* | 17.438 | OR020791 | OR051968 | OR020649 | OR637276 |  |  |  |
| Gasterosteidae | *Gasterosteus* | *aculeatus* | 16.609 | OR020792 | OR051969 | OR020650 | OR637277 |  |  |  |
| Gasterosteidae | *Pungitius* | *pungitius* | 16.583 | OR020793 | OR051970 | OR020651 | OR637278 |  |  |  |
| Hiodontidae | *Hiodon* | *alosoides* | 16.648 | OR020794 | OR051971 | OR020652 | OR637279 |  |  |  |
| Ictaluridae | *Ameiurus* | *melas* | 16.513 | OR020795 | OR051972 | OR020653 | OR637280 |  |  |  |
| Ictaluridae | *Ameiurus* | *natalis* | 16.512 | OR020796 | OR051973 | OR020654 | OR637281 |  |  |  |
| Ictaluridae | *Ameiurus* | *nebulosus* | 16.513 | OR020797 | OR051974 |  | OR637282 |  |  |  |
| Lotidae | *Lota* | *lota* | 16.562 | OR020798 | OR051975 | OR020655 | OR637283 |  |  |  |
| Osmeridae | *Osmerus* | *mordax* | 16.660 | OR020799 | OR051976 | OR020656 | OR637284 |  |  |  |
| Osmeridae | *Spirinchus* | *thaleichthys* | N/A | OR020800 | OR051977 | OR020657 | N/A |  |  |  |
| Osmeridae | *Thaleichthys* | *pacificus* | 16.742 | OR020801 | OR051978 | OR020658 | OR637285 |  |  |  |
| Percidae | *Perca* | *flavescens* | 16.668 | OR020802 | OR051979 | OR020659 | OR637286 |  |  |  |
| Percidae | *Sander* | *vitreus* | 16.884 | OR020803 | OR051980 | OR020660 | OR637287 |  |  |  |
| Percopsidae | *Percopsis* | *omiscomaycus* | 16.710 | OR020804 | OR051981 | OR020661 | OR637288 |  |  |  |
| Petromyzontidae | *Entosphenus* | *macrostomus* | 16.585 | OR020805 | OR051982 | OR020662 | OR637289 |  |  |  |
| Petromyzontidae | *Entosphenus* | *tridentatus* | 16.124 | OR020806 | OR051983 | OR020663 | OR637290 |  |  |  |
| Petromyzontidae | *Lampetra* | *ayresii* | 16.695 | OR020807 | OR051984 | OR020664 | OR637291 |  |  |  |
| Petromyzontidae | *Lampetra* | *richardsoni* | 16.150 | OR020808 | OR051985 | OR020665 | OR637292 |  |  |  |
| Petromyzontidae | *Lampetra* | *richardsoni (Morrison Creek)* | 16.225 | OR020809 | OR051986 | OR020666 | OR637293 |  |  |  |
| Pleuronectidae | *Platichthys* | *stellatus* | 16.985 | OR020810 |  | OR020667 | OR637294 |  |  |  |
| Salmonidae | *Salvelinus* | *confluentus (Coastal)* | N/A | OR020811 | OR051987 | OR020668 | N/A |  |  |  |
| Salmonidae | *Salvelinus* | *confluentus (Interior)* | N/A | OR020812 | OR051988 | OR020669 | OR637311 |  |  |  |
| Salmonidae | *Coregonus* | *artedi* | 16.538 | OR020813 | OR051989 | OR020670 | OR637295 |  |  |  |
| Salmonidae | *Coregonus* | *clupeaformis* | 16.739 | OR020814 | OR051990 |  | OR637296 |  |  |  |
| Salmonidae | *Coregonus* | *nasus* | 16.537 | OR020815 | OR051991 | OR020671 | OR637297 |  |  |  |
| Salmonidae | *Coregonus* | *sardinella* | 16.529 | OR020816 | OR051992 | OR020672 | OR637298 |  |  |  |
| Salmonidae | *Oncorhynchus* | *clarki* | 16.658 | OR020817 | OR051993 | OR020673 | OR637299 |  |  |  |
| Salmonidae | *Oncorhynchus* | *clarki (Lewisi)* | 16.696 | OR020818 | OR051994 |  | OR637300 |  |  |  |
| Salmonidae | *Oncorhynchus* | *gorbuscha* | 16.657 | OR020819 | OR051995 |  | OR637301 |  |  |  |
| Salmonidae | *Oncorhynchus* | *keta* | 16.655 | OR020820 | OR051996 | OR020674 | OR637302 |  |  |  |
| Salmonidae | *Oncorhynchus* | *kisutch* | 16.660 | OR020821 | OR051997 | OR020675 | OR637303 |  |  |  |
| Salmonidae | *Oncorhynchus* | *mykiss* | 16.660 | OR020822 | OR051998 | OR020676 | OR637304 |  |  |  |
| Salmonidae | *Oncorhynchus* | *nerka* | 16.634 | OR020823 | OR051999 | OR020677 | OR637305 |  |  |  |
| Salmonidae | *Oncorhynchus* | *tshawytscha* | 16.540 | OR020824 | OR052000 | OR020678 | OR637306 |  |  |  |
| Salmonidae | *Prosopium* | *coulterii* | 16.772 | OR020825 | OR052001 | OR020679 | OR637307 |  |  |  |
| Salmonidae | *Prosopium* | *cylindraceum* | 16.912 | OR020826 | OR052002 | OR020680 | OR637308 |  |  |  |
| Salmonidae | *Prosopium* | *williamsoni* | 16.754 | OR020827 | OR052003 | OR020681 | OR637309 |  |  |  |
| Salmonidae | *Salmo* | *salar* | 16.672 | OR020828 | OR052004 | OR020682 | OR637310 |  |  |  |
| Salmonidae | *Salvelinus* | *fontinalis* | 16.619 | OR020829 | OR052005 | OR020683 | OR637312 |  |  |  |
| Salmonidae | *Salvelinus* | *malma* | 16.651 | OR020830 | OR052006 | OR020684 | OR637313 |  |  |  |
| Salmonidae | *Salvelinus* | *namaycush* | N/A | OR020831 | OR052007 | OR020685 | OR637314 |  |  |  |
| Salmonidae | *Stenodus* | *leucichthys* | 16.845 | OR020832 | OR052008 | OR020686 | OR637315 |  |  |  |
| Salmonidae | *Thymallus* | *arcticus* | 16.661 | OR020833 | OR052009 | OR020687 | OR637316 |  |  |  |
