## Supplementary Table 3 for "NAMERS: a purpose-built reference DNA sequence database to support applied eDNA metabarcoding"

|  | Catostomidae | Cyprinidae | Osmeridae | Salmonidae |
| --- | --- | --- | --- | --- |
|  | *Pantosteus* | *Margariscus* | *Spirinchus* | *Salvelinus* |
|  | *bondi* | *nachtriebi* | *thaleichthys* | *confluentus (Coastal)* |
| COX1 |  | OR557495 | OR557496 | OR557497 |
| COX2 | OR567801 | OR567800 | OR567802 |  |
| COX3 | OR567803 |  | OR567804 |  |
| CYTB | OR567805 |  | OR567806 | OR567807 |
| ND1 | OR567808 |  | OR567809 |  |
| ND2 |  |  | OR567810 |  |
| ND3 | OR567812 | OR567811 | OR567813 | OR567814 |
| ND4 | OR567815 |  | OR567816 | OR567817 |
| ND4L |  |  |  | OR567818 |
| ND5 | OR567820 | OR567819 | OR567821 |  |
| ND6 |  |  | OR567822 |  |
| ATP6 | OR567823 |  | OR567824 |  |
| ATP8 | OR567825 |  | OR567826 |  |
| s-rRNA | OR559979 | OR559978 | OR559980 | OR559981 |
| l-rRNA | OR557499 | OR557498 | OR557500 |  |
| 5.8S | OR020755 |  | OR020800 | OR020811 |
| 18S | OR051933 | OR051957 | OR051977 | OR051987 |
| 28S | OR020615 |  | OR020657 | OR020668 |
